# The ecology of remembering and forgetting: quantity-quality trade-offs in the spatiotemporal memory of a forager

**DOI:** 10.64898/2026.07.12.738111

**Authors:** Kavel C. D. Ozturk, Bryndan O. C. M. van Pinxteren, Karline R. L. Janmaat, Benjamin Robira

## Abstract

One way that animals can cope with the challenge of locating ephemeral food in time and space is by tracking elapsed time and planning revisits to these resources. These abilities likely evolved under constraints of memory capacity. We expect that during evolution a trade-off emerged between a large memory size (quantity) and high accuracy (quality) of information. We used computer simulations of a forager moving through space and time to investigate how timing accuracy, memory size, and forgetting affect foraging efficiency across environmental conditions, and how foragers should trade off the quantity against quality of memorised information. Memory in general paid off, as it improved foraging efficiency. However, surprisingly, the largest accuracy and size were not always most beneficial. In resource-poor, heterogeneous, and highly dynamic environments, extensive memory was even detrimental, as individuals likely became trapped in overexploited familiar areas. This suggests that under certain environmental conditions, a hidden, non-energetic cost of memory can arise. Furthermore, environmental structure shaped a quantity–quality trade-off such that a minimal memory size and higher timing accuracy were favoured in resource-poor, temporally stable and homogenous environments. Finally, forgetting was beneficial when memory was constrained, and environments were poor, heterogenous and dynamic. Forgetting limited the benefits of increased memory size, highlighting that memory costs can emerge from how it shapes movement patterns and foraging decisions. Overall, our results highlight that larger and more accurate memory is not necessarily better and that forgetting can be adaptive. This study takes a first step toward the theoretical consideration of memory trade-offs in order to research how they shape, and are shaped, by foraging pressures.

**Author summary:** How can we explain foraging memory abilities differences in animals? In this study, we make an attempt to elucidate the current patterns of temporal memory using a computational model that was inspired by realistic environmental and cognitive mechanisms. Building on our previous theoretical and empirical investigations of the causes and consequences of spatiotemporal memory, we provide new insights into the ecological drivers of temporal memory, how they shape an accuracy-size trade off, and its consequences for movement behaviour. We show that a larger memory is favoured, at the cost of accuracy, in resource-rich and dynamic environments. Our results highlight that while memory is generally beneficial, a larger and more accurate memory is not necessarily better. Moreover, our model suggests that such constraints of memory stem from a hidden cost of how memory restricts movement and instigates local overexploitation. Our findings contribute to a broader understanding of how cognition evolves in response to ecological conditions.

## 1. Introduction

For a mobile forager, finding food can be a challenge as it often requires the ability to discern *when* and *where* to look for food. A plethora of animal species have been shown to be able to remember the locations of a multitude of food sources in their environment and to use various memory-based strategies to travel to these locations (see [1] for a review of primate foraging decisions and cognitive mechanisms; and [2] for a review of the recollection of past events and planning in animals). For example, various mammal and bird species use memory to plan revisits to ephemeral food sources based on the elapsed time and the renewal, degradation or depletion of these food sources [3–13]. These findings raise the question of how animals evolved to remember the passage of time during foraging; How did this ability affect foraging efficiency and how has temporal memory been shaped by selection?

When an animal searches for an ephemeral and scattered resource, it can move in a straight line towards remembered food sources, avoiding costly detours [1, 14]. Yet, to be the most efficient, it should not only remember resource locations, but also know which ones currently contain food [15, 16]. *En route* to the target, or at the target itself, a forager can thus collect information on the state of availability of resources that may later be used to infer availability (in time) [16]. For example, chimpanzees (*Pan troglodytes*) have been observed to monitor 87% of their potential food trees for ripe fruit availability on their way to other food trees [14]. This monitoring behaviour can provide an individual with temporal information about when an event (e.g., when a tree with unripe fruits was encountered) occurred in multiple ways. One way that a remembered event can be placed in time is by directly tracking how much time has elapsed since the event (also called distance-based or relative timing) [17, 18]. Alternatively, when an event occurred can be reconstructed indirectly from contextual information stored in memory (such as the location of the tree, what other food sources were available; [17, 18]). In any case, a common denominator of any temporal memory type is some notion of elapsed time, either by tracking, or reconstructing, elapsed time.

The ability to track time is prone to errors, which may induce discrepancy between the “true” information and the “remembered” one [19, 20]. These errors may clearly impact the benefits of temporal memory. For example, ripe fruits - an important and high-energy food source for a variety of animals - on individual trees can be largely depleted within a time frame of several days in tropical forests [21, 22], hence rapidly leading to outdated information. Furthermore, this food source can be extremely scarce in tropical forests, with an encounter rate of one large ripe fruit crop per 21 km of straight travel in some tropical forests [15], thus creating a narrow time window for finding it and making errors in time tracking very costly.

Errors in memories can arise due to processes during memorisation itself (constructive errors; [23, 24]), or they can happen later during memory recall (reconstructive errors; [23, 25]). In the latter, the inaccuracy may depend on dynamic and fixed traits. Dynamic traits may be linked to the individual (e.g., age; [26]) but can also be related to the elapsed time itself, where the quality of time estimation is associated with the elapsed time. For example, individuals tend to date older events less accurate [18, 27]. In addition to this general inaccuracy, there is a tendency to consider past events as older than they really are, and recent events as more recent than they really are, a bias known as telescopic effect [28].

In parallel, the fixed cognitive machinery may itself induce errors. In the intertwining of memories, confusion may occur through interference, as memories compete for access during recall [29, 30]. In other words, the accuracy of recalling may be constrained by the ability of the cognitive machinery to process and sort out a certain amount of information without error. As such, a quantity-quality trade-off may emerge [31], where individuals memorise a large amount of information with poor accuracy, or little information with high accuracy. This relationship between the number of memories (quantity) and recall accuracy (quality) may not simply be linear (i.e., a linear decrease in accuracy with increasing memory size), but may grow non-linearly, as observed, for example, in visual working memory [32].

Beyond being prone to error, extended memory also comes with some costs [31] bringing about two major consequences. Firstly, memory capacity (i.e. the amount of information that can be retained in memory) must be limited [33]. This limitation likely renders mechanisms such as forgetting not a simple defect of memory, but an adaptive feature that enables a dynamic adjustment of the knowledge to current conditions (e.g., it is likely beneficial to forget outdated memories [34]). Specifically, mechanisms of active forgetting (*versus* passive forgetting), where specific memories are erased (or their accessibility is inhibited) could play such an adaptive role in a foraging context [35–42]. Secondly, memory capacity limitations may be subject to selection itself [43]. Different environments may render memory use more or less beneficial [43], hence what quality and quantity of memories combined should be evolutionarily favoured.

It has been suggested that environments with intermediate levels of spatiotemporal complexity should be most favourable for spatiotemporal memory [16, 31, 44]. In ephemeral, spatially heterogenous, and low-productive environments, a forager could benefit significantly from remembering and returning to potential resources that might otherwise be difficult to locate in both space and time. For example, when environments are temporally stable, spatially homogenous and highly productive, tracking elapsed time to return to remembered trees that previously had unripe fruits would have little added value, as resources are highly predictable and could be located by use of sensory cues. In such environments, an extensive temporal memory capacity could lead to local resource depletion and missed foraging opportunities elsewhere. Conversely, when resources are extremely ephemeral, heterogenous and scarce, up-to-date information may be so limited that memory provides little benefit. Finally, for a given environment, the memory capacity limitation is expected to shape the quantity-quality trade-off. Yet we still lack a theoretical understanding of how environmental spatiotemporal heterogeneity should shape this trade-off to maximise foraging efficiency.

In this study, we theoretically tested how the quantity-quality trade-off is expected to vary with environmental heterogeneity to maximize foraging efficiency. We used an agent-based model (ABM) simulating the movement and foraging behaviour of an artificial animal (hereafter: forager) roaming an environment composed of ephemeral resources. This forager was cognitively endowed (i.e. a memory-guided forager). However, when it remembered where and when to find (some of) the resource patches it had a certain inaccuracy and a forgetting mechanism. Specifically, we considered variations in memory size (i.e. the “quantity” feature), memory inaccuracy (timing error; i.e. the “quality” feature) and forgetting mechanism (i.e. active versus passive forgetting). We then compared the forager’s efficiency between environments differing in their spatial and temporal heterogeneity. To deepen our understanding of the foraging benefits provided by spatiotemporal memory, we considered these variations relative to those expected when a forager was only capable of short-range detection of food sources in the environment (i.e. a naive forager). Our model was inspired by biologically realistic environmental and cognitive models (of <u>tracking elapsed time</u>: [10, 11, 13, 45]; <u>forgetting mechanisms</u>: [35, 38–41, 46]), making it possible to assess the adaptive value of spatiotemporal memory in animal foragers overall.

## 2. Material & Methods

### 2.1 Agent-Based Model Philosophy

Based on a minimalistic set of assumptions, we aimed to test (1) whether a simulated forager’s efficiency is more strongly enhanced by increased quality (i.e., reduced timing error) and/or quantity (i.e., size) of memory of potential resources, (2) how the quality-quantity trade-off is shaped by resource spatiotemporal patterns, and (3) how forgetting mechanisms affect memory benefits and this trade-off. For this, we used *R* software (v.4.1.2)[47] to perform agent-based simulations, analyses, and visualisations.

For simplicity, and based on our expertise, the virtual system was initially ideated to resemble a frugivore primate looking for ripe fruit-bearing trees in a tropical forest (as in [16, 48]). Specifically, we simulated the behaviour of a single solitary forager (hereafter forager; see **Movement rules**) in a set of environments that varied in resource temporal availability, density and spatial distribution. This non-competitive context facilitates computational simulation execution and interpretation, while ensuring biological realism (since in competitive environments memory should also drive the spatial segregation of individuals [49]).

Since all spatial and temporal units in our model are arbitrary and we cover a large palette of cognitive abilities and resource distributions, the model can, depending on the parameterisation, serve as a simplified representation of many natural systems. This makes the conclusions of this theoretical model largely generalisable. The parameterisation of this model is available in S1 Table.

To answer (1) how advantageous a large/accurate temporal memory is, and (2) how the quality-quantity memory trade-off is shaped by environmental conditions, we tested different foraging conditions (see *section* 2.4 **Foraging scenarios and analysis**). Here we varied (a) the forager’s cognitive abilities in terms of memory accuracy and size, and an associated interference costs whereby increasing memory quantity reduces memory quality.

Furthermore, we varied (b) the spatial and temporal distribution of resources and measured the variations in forager’s performance, quantified as the foraging efficiency (i.e., the amount of food collected within each simulation).

### 2.2 Environment

#### 2.2.1 Resource spatial distribution

The environment was modelled as a two-dimensional flat square area of 1000 x 1000 arbitrary-length units (alu) (See S1 Table). This environment included 25 to 3200 resource patches – hereafter, (fruit) trees – that were either homogeneously or heterogeneously distributed. We simulated homogeneous environments by sampling each tree location’s *x* and *y* coordinates from a uniform distribution (from 0 to 1000). We simulated spatially heterogeneous environments by sampling tree locations from ten circular normal distributions whose centre locations’ coordinates were, again, sampled from a uniform distribution. The degree of spatial heterogeneity was altered by varying the standard deviation, defined as the tree patch spread, of the circular normal distribution. Spatial heterogeneity values (i.e. the standard deviation of the circular normal distribution) ranged from five arbitrary length units (resulting in low patch spread, hence a small patch size) to 320 alu (resulting in high patch spread, corresponding, hence a larger patch size; Fig 1A; S1 Table). These distributions were determined and fixed at the beginning of each simulation, resulting in a spatially fixed environment.

**Fig 1.**
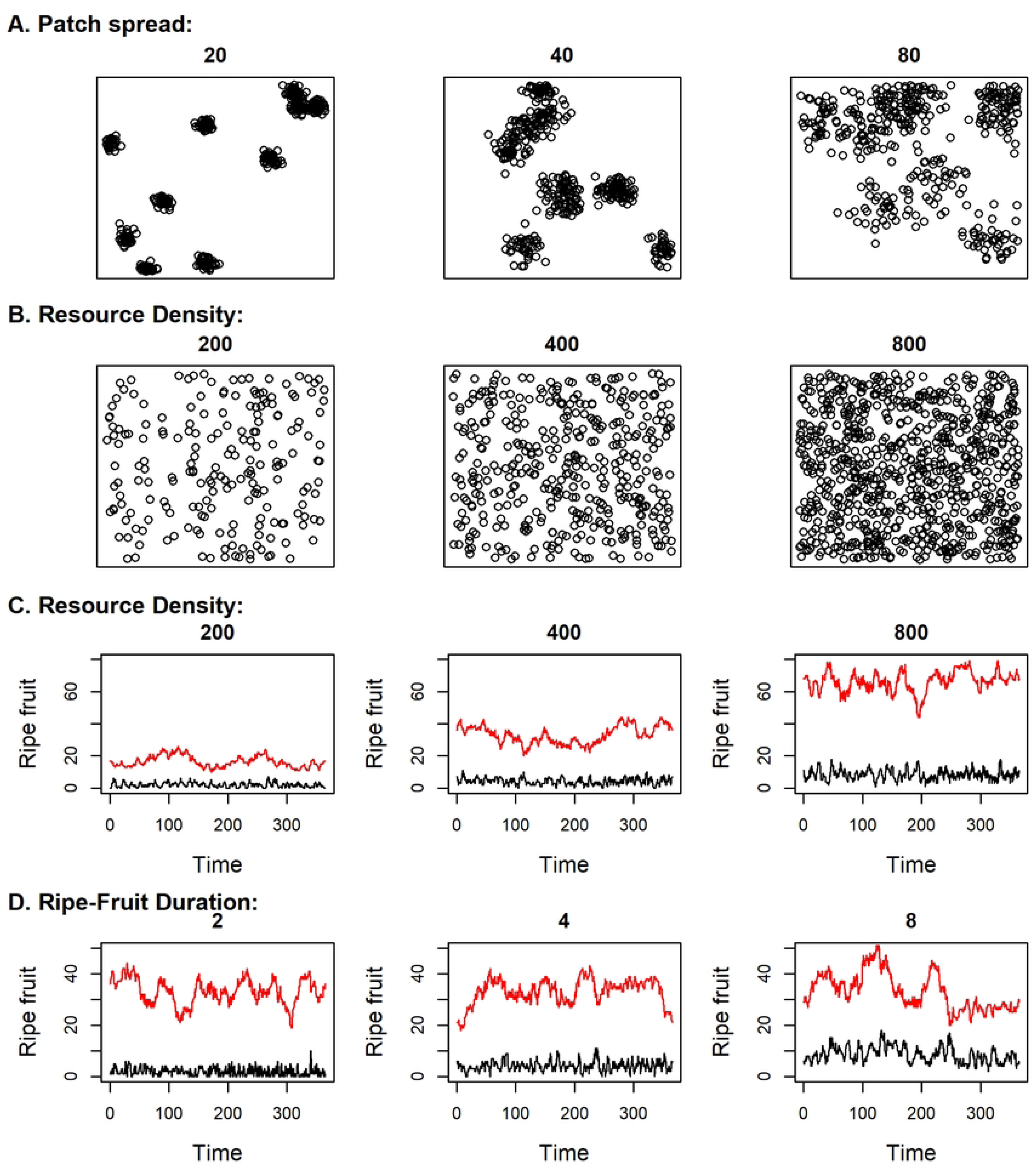
Aerial view of the spatial resource distribution and time series of fruit abundance. (A & B) Spatial resource distribution and time series (C & D) of ripe (black) and unripe (red) fruit abundance in the simulated environments with different values for patch spread (A), resource density (B & C) and ripe-fruit duration (D). The values for patch spread, resource density and ripe fruit duration are indicated in their respective plots. The parameters that were not indicated in the plot were set to their default values (See S1 Table).

#### 2.2.2 Resource temporal distribution

We considered each tree to belong to the same species. Each tree produced one arbitrary fruit unit (afu), with a constant interval of 365 time units (tu), mimicking a “year”. The start of the first fruit production for each tree was randomly drawn from a uniform distribution over a year, resulting in asynchronous fruit production (such as is the case for various fig species that form an important food source in tropical ecosystems [50]) throughout the environment. The one afu appeared immediately at the start of the fruiting period, but only became ripe and edible after 30 tu (as with some unripe fruits that are less palatable or accessible due to chemical or mechanical protection when unripe [51]). After ripening, the one afu persisted for a tu-range of 0.25–31 (depending on the simulated environmental conditions), unless it was depleted earlier by the forager. By varying this ripe-fruit availability period (hereafter: ripe-fruit duration), we modelled variation in the strength of processes such as fruit degradation and depletion.

### 2.3 Movement rules

At the start of the simulation, a single forager started in the middle of the simulated environment. This forager would move with a speed of 500 alu.tu^−1^ following three movement mechanics selected hierarchically in this order: sensory-based movement, memory-based movement, or random movement. Specifically, at each simulation step, the forager acted based on a fixed priority sequence (Fig 2). For this it 1) attempted to eat food at its current location, and if possible it would eat at a speed of 1 afu per 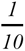 tu; failing that, it 2) moved towards detectable food (sensory-based movement); if no food was detectable, it 3) moved towards remembered food, if any (memory-based movement); and if none of these options were possible, it 4) moved randomly.

**Fig 2.**
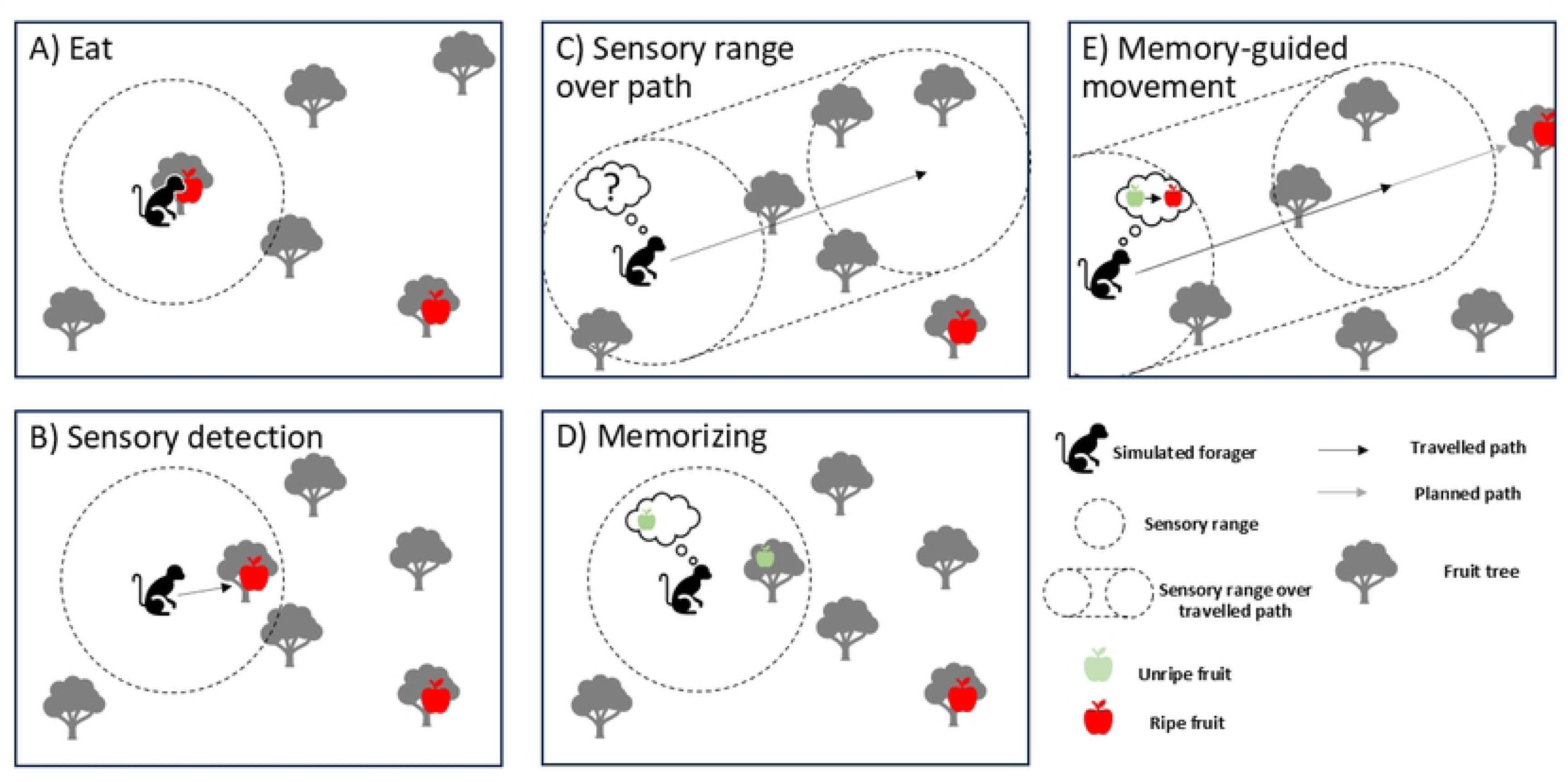
Overview of the major foraging rules that affect the forager’s movement. When the simulated forager is at a tree with ripe fruit it will eat (A). If the forager is not eating, it will move towards a tree with ripe fruit, if there is one in its sensory range (B). When the forager is not eating and when no food could be sensed (or when it has no memories that it expects to contain ripe fruit) the forager will move in a random fashion (C). If trees with unripe fruits come within the sensory range over the movement path the forager will store the locations and the time the tree will have ripe fruits in its memory (D). When the forager is not eating and there is no ripe fruit within its sensory range, the forager may assess whether it expects an out-of-sight remembered tree to contain ripe fruit. If the forager expects there to be ripe fruit at a remembered tree, it will move towards the remembered tree (E). If none of the other cases are true, the forager will move randomly.

#### 2.3.1 Sensory-based movement

The forager was aware of all trees within its sensory range and would move to the closest detected tree with ripe and edible fruit, if any was present. This sensory range, *d*, depended on the tree density and was set to 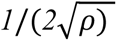 where ρ is the tree density of the environment [52]. This corresponds to the expected nearest-neighbour distance under a completely homogeneous distribution. This sensory range was selected to ensure a balanced contribution of sensory and memory-based processes to movement.

#### 2.3.2 Memory-based movement

The memory of the forager was defined by its accuracy, but also by its maximum memory size, both of which could vary between simulations (see **2.4 Foraging scenarios and analysis**; S1 Table). When using memory (see **2.4 Foraging scenarios and analysis**), the forager had the ability to remember (without a prior learning process) with perfect accuracy both the locations of trees that contained unripe fruits at past encounters and the date at which trees would be expected to contain ripe fruit. However, the forager had no knowledge of the fruiting duration and inaccurately kept track of elapsed time. Specifically, for a tree *i* that was encountered at time *t* with unripe fruit, the forager pinpointed the time, *T_i_*, it would take for fruit to appear. As the forager performed a movement step from time *t+μ to t+μ’,* where *μ* and *μ’* are the true time elapsed since the tree was last encountered at the beginning and end of the step, respectively, it updated its estimated elapsed time from *T_μ_* to *T_μ_*_’_. At time *t+μ’,* the forager estimates that *T_μ_*_’_ time units have elapsed, where *T_μ_*_’_ is updated according to *T_μ’_* =*T_μ_* + TN(*μ’ - μ, λ; a =* -*T_μ_*, *b =* ∞), where *λ* represents the <u>timing error</u> parameter and TN(*m, sd; a, b*) denotes a truncated normal (Gaussian) random variable with mean *m* and standard deviation *sd*, truncated to the interval [*a*, *b*]. Due to the truncation of the Gaussian distribution in its lower bound, the estimate of elapsed time does not take negative values (i.e. it never extends into the future). Together with the temporal autocorrelation of the estimates, this truncation induces a slight backward telescoping effect (S1 Fig), where the estimated elapsed time exceeds the actual elapsed time on average as has been found in various studies [18, 19].

Based on the memory of tree locations and elapsed time, the forager could then engage in memory-based movement towards the tree for which the predicted fruiting would be the nearest in estimated time (i.e., for which *Δt* = *T_i_ - T_μ_*_’_) while accounting for the time it would take to reach resources (travel time τ*_i_*). The probability for the forager to target this location (or otherwise engage in random movement) was defined as

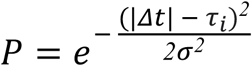

where σ is a scale parameter.

When, during the current time step or before, a memorized tree had been targeted, the forager would move by consecutive steps of 100 alu in a straight line towards the memorized location of the targeted tree, analogous to behaviour observed in the wild [14]. However, the forager’s route could be adjusted based on encountered fruit trees along the way. For each movement step, if the forager encountered ripe fruit along its path, it would move from the first point of detection straight towards the first detected tree with ripe fruit on the route.

#### 2.3.3 Learning and Forgetting

Within a simulation, the forager’s memory was dynamic, incorporating both learning and forgetting. When the forager moved and detected one or more previously unknown trees bearing unripe fruits, it memorized (in order of detection) each tree’s location and predicted fruit ripening date (based on the maturation period), provided that memory space was available (Fig 2D). In case a memorized tree with unripe fruit was detected, the time elapsed since encounter was reset to 0.

At each movement step, the probability for the retention of a memorized tree in memory was defined by a simple power law: *R = t*^-0.001^, where *t* is the actual elapsed time since the tree is memorized [46, 53]. Additionally, a memorized tree would be discarded from the forager’s memory if the tree was detected without fruit, i.e. a process of actively forgetting. (While we call it *active* forgetting by opposition to the *passive* continuous fading of old memories, this does not mean that such a process needs to be conscious.) In the “only passive forgetting” scenario (see **2.4 Foraging scenarios and analysis**), the process of active forgetting was omitted.

#### 2.3.4 Random movement

When no sensory or memorized information (that could be acted upon) was available, the forager moved randomly. This random movement corresponded to a correlated random walk (CRW) of 100 alu in which there is a correlation between the direction of movement between two successive steps, in order to capture natural advection traditionally expected in animal movement [54]. To do so, we drew the change in direction between consecutive steps, i.e. the turning angle, from a zero-centred wrapped Normal distribution with a concentration parameter of 0.9. This value corresponds to strongly but not perfectly directed movement (angular standard deviation ≈ 26° of the unwrapped normal distribution, i.e. most turns within roughly ±50° of the previous heading). This produces moderately tortuous paths consistent with advective movement rather than pure diffusion. If the agent had not moved during the previous time step (because the simulation just started, or due to the forager just having eaten), the direction of the random movement was drawn from a uniform distribution. If the forager reached an environmental boundary during the CRW, it was reoriented from the collision point in a random direction that kept the forager in the environment for the remainder of the movement. During the CRW, the forager would adjust movement if it perceived a tree with ripe fruit along its path. In such cases, the forager would move from the first point of detection straight towards the first detected tree with ripe fruit on the route.

### 2.4 Foraging scenarios and analysis

Each simulation was preceded by an initialization phase of 60 tu, in order to allow the forager to create memories. We then examined the foraging efficiency *E* of the forager only in the operation phase, which had the length of one fruiting interval, i.e. (365 tu) and quantified it as the number of *afu* consumed. We considered three scenarios:

#### 2.4.1 Scenario 1. How advantageous is memory, and when?

To understand the overall benefit of memorising a large amount of information and the precision of memorising, we first simulated foragers endowed with various memory sizes (from 1 to 25 known trees, with an increment of two), and memory accuracies (*λ* varying from 0.01 to 1 with an increment of 0.09). We then considered 24 environmental settings which varied along one parameter at a time, namely ripe-fruit duration, resource density or patch spread (each considering 8 cases). When we varied one environmental parameter, the other two were fixed at their reference value (Table S1). Additionally, in the Supporting Information (Appendix A in S1 File), we calculated the core area of the simulated agents in Scenario 1 for various environmental parameters.

#### 2.4.2 Scenario 2. Should you know more or better?

In the second scenario, we investigated the optimality of the quality-quantity trade-off. For this, we used the parameter and foraging efficiency values of scenario 1, to examine how a <u>non-dynamic</u> linear trade-off between quality and quantity is shaped by the environment. (But, see Appendix B in S1 File, for a dynamic trade-off between memorized temporal information where information is less accurate due to interference from other memories that were retained at the same time.) This relationship was chosen such that the inaccuracy parameter *λ* (which was constant during each simulation) was linearly related to the maximum number of memories, *λ =* 0.99/22*N* - 0.77/22, with *N* representing the maximum number of memories that a forager was able to hold.

#### 2.4.3 Scenario 3. Should you forget or not?

To understand the role of active forgetting (i.e. the active removal of specific memorized information) on memory benefits, we modelled a memory-guided forager that was identical to the foragers in scenario 1 except for the ability of active forgetting. A memory-guided forager in scenario 3 would not actively forget memorized empty trees that it encountered. Instead, the forager would retain the fruitless tree in its memory and infer that it would fruit in the distant future (i.e. outside of the simulated time). Thereby, this tree could only be forgotten due the passive forgetting process.

#### 2.4.4 Foraging Efficiency

For scenario 1 and 2, we considered two types of foragers: a forager that could only rely on detection but had no memory (naive forager), and a forager that relied on both sensory detection and memory (memory-guided forager). The naive forager would highlight the expected baseline foraging efficiency, which can vary depending on the environment spatio-temporal pattern [16]. As such, we defined the relative foraging efficiency *E_r_* as 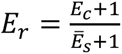, where *E_c_* represents the absolute foraging efficiency of the memory-guided forager and *E̅_s_* was the average foraging efficiency of the naive foragers with the same environmental parameters. We added 1 to both the numerator and denominator to make the relative efficiency less sensitive in scenarios where the efficiency of naive foragers is very low or near zero. We then used *E_r_* as a proxy of fitness to discuss how the trade-off quality *vs* quantity was shaped when the environment varied in terms of resource density, ripe-fruit duration, and patch spread.

For scenario 3, similar to scenario 1 and 2, we determined the relative foraging efficiency, *E_r_* of memory-guided foragers without active forgetting abilities, as 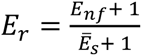. Here, *E_nf_* was the absolute foraging efficiency of the foragers without active forgetting abilities. We conducted 300 simulations for each resulting combination of environmental and cognitive parameter values.

## 3. RESULTS

### 3.1 Scenario 1: Memory is advantageous, but the magnitude of the advantage varies greatly with resource spatiotemporal distribution

When accurate and extensive knowledge was at no cost, memory-based foraging increased efficiency, but not uniformly. We found that the relative foraging benefit varied substantially across different memory and environmental parameters. Memory-guided foraging was most beneficial when 1) the ripe-fruit duration was short, and 2) environments were resource-poor and high to moderately heterogeneous. With short ripe-fruit duration (0.25 tu), the best (i.e., across all possible quantity-quality combinations) memory-based forager achieved up to a 7.3-fold higher efficiency than a naive forager (Fig 3; Fig 4A-C). Similarly, in environments low in resource density (25 × 10⁻⁶ trees/alu²), memory increased efficiency by up to 2.1-fold (Fig 3; Fig 4D-F). Finally, under high (2.5 alu) and moderate (20 alu) patch spread, memory-based foraging was most strongly favoured, yielding a higher advantage compared with a reliance on sensory cues only (up to a 2.1 and 1.6-fold advantage with and without active forgetting, respectively) (Fig 3; Fig 4G-I). On the other hand, in environments with long ripe-fruit duration, moderate to high resource density, or high patch spread, memory-guided foraging often provided little or no advantage. In some cases, these foragers even performed worse than naive foragers, resulting in negative relative benefits (Fig 4).

**Fig 3.**
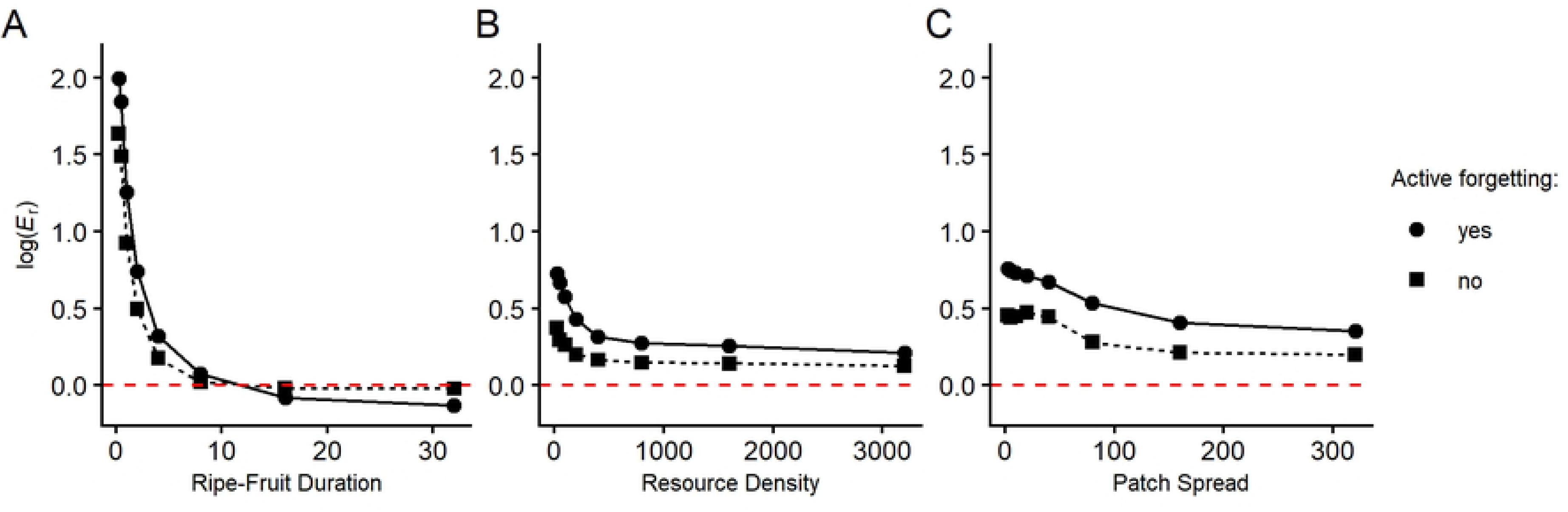
Memory-guided foraging is most beneficial when the ripe-fruit duration is short, environments are resource-poor and moderately heterogeneous. Relative foraging efficiency (log-transformed) of the best-performing memory-guided foragers of the two types of foragers (with and without an active forgetting mechanism) across environments that varied in ripe-fruit duration (A), resource density (B) and patch spread (C). Points correspond to the average transformed relative foraging benefit across 300 simulations.

**Fig 4.**
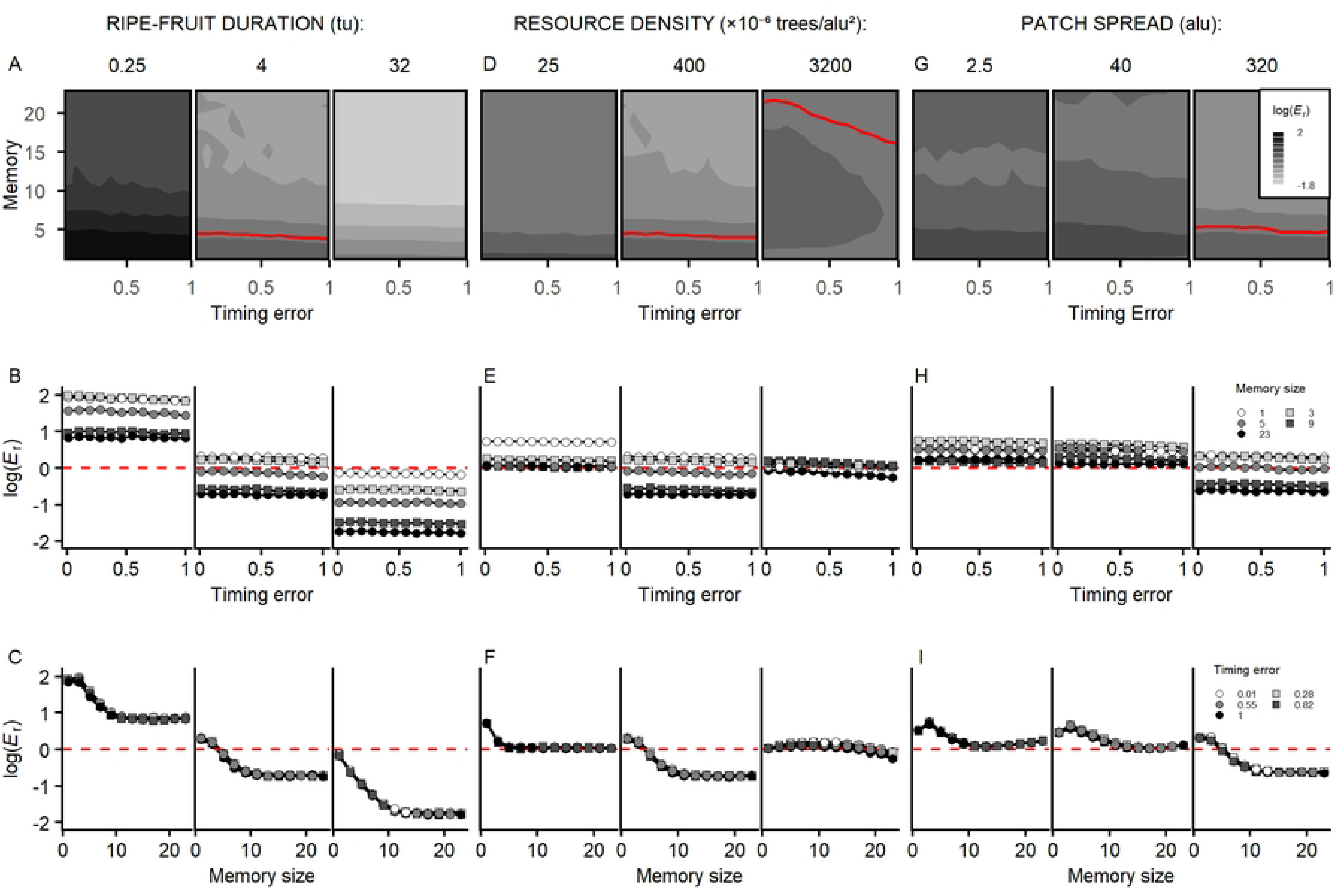
Ripe-fruit duration, resource density and patch spread shape the benefits of memory size, but not timing accuracy. Relative foraging efficiency (log-transformed) of the memory-guided foragers that differed in their cognitive abilities (timing error and memory size) for various values of ripe-fruit duration (A-C), resource density (D-F) and patch spread (G-I). The plots in the second row (B, E, H) and third row (C, F, I) show cross-sections of the contour plots (A, D, G) for different values of memory size and timing error, respectively. The red lines indicate the threshold where there is no relative foraging benefit compared to a naive forager.

The foraging benefits were shaped by the environmental conditions, but also depended on the quantity and quality of the temporal memory abilities. Improvements in timing accuracy generally increased the foraging benefit, although the effect was modest (with the relative efficiency improving 20.7% when comparing foragers with a minimal timing error versus maximal timing error). With small-to-intermediate memory size the effect of timing accuracy was more pronounced (S2 Fig). Yet, with large memory size, timing accuracy exerted a smaller influence. Low-to-intermediate levels of memory size mostly maximized the relative foraging benefit (Fig 4C; 4F; 4I), with initial increases improving foraging benefit, but further increases in memory size resulting in a decline. However, this optimal efficiency shifted toward a smaller memory size when resource density was low and ripe-fruit duration was long, and toward a larger memory size when resource density was high (Fig. 4C, 4F).

Additionally, the area of the environment that was predominantly used by the forager decreased with increasing memory size (S3 Fig). As such, the best-performing memory parameters were not simply those with a combination of the highest accuracy and the largest memory size (S4 Fig). Instead, intermediate memory size paired with high (but not always maximal) timing accuracy performed best, with the environmental conditions shaping the optimal balance between quality and quantity of memory.

### 3.2 Scenario 2: The most accurate or largest memory size does not necessarily lead to the highest foraging efficiency

In scenario 2, we introduced a fixed trade-off between the quality and quantity of memories (where the cost of memory is determined by the maximum memory size). When introducing a quality-quantity trade-off, the optimal memory size changed substantially with changes in resource density (Fig 5; Fig 6). In environments with a low resource density, foragers benefited most from minimal memory size and thus maximal timing accuracy. As resource density increased, the optimum shifted toward a larger memory size and lower timing accuracy. Finally, in resource-dense environments, an intermediate level of memory size was optimal. The optimal memory size shifted more gradually with changes in patch spread and ripe-fruit duration, towards a larger memory size with a shortening ripe-fruit duration and decreasing patch spread (Fig 5; Fig 6).

**Fig 5.**
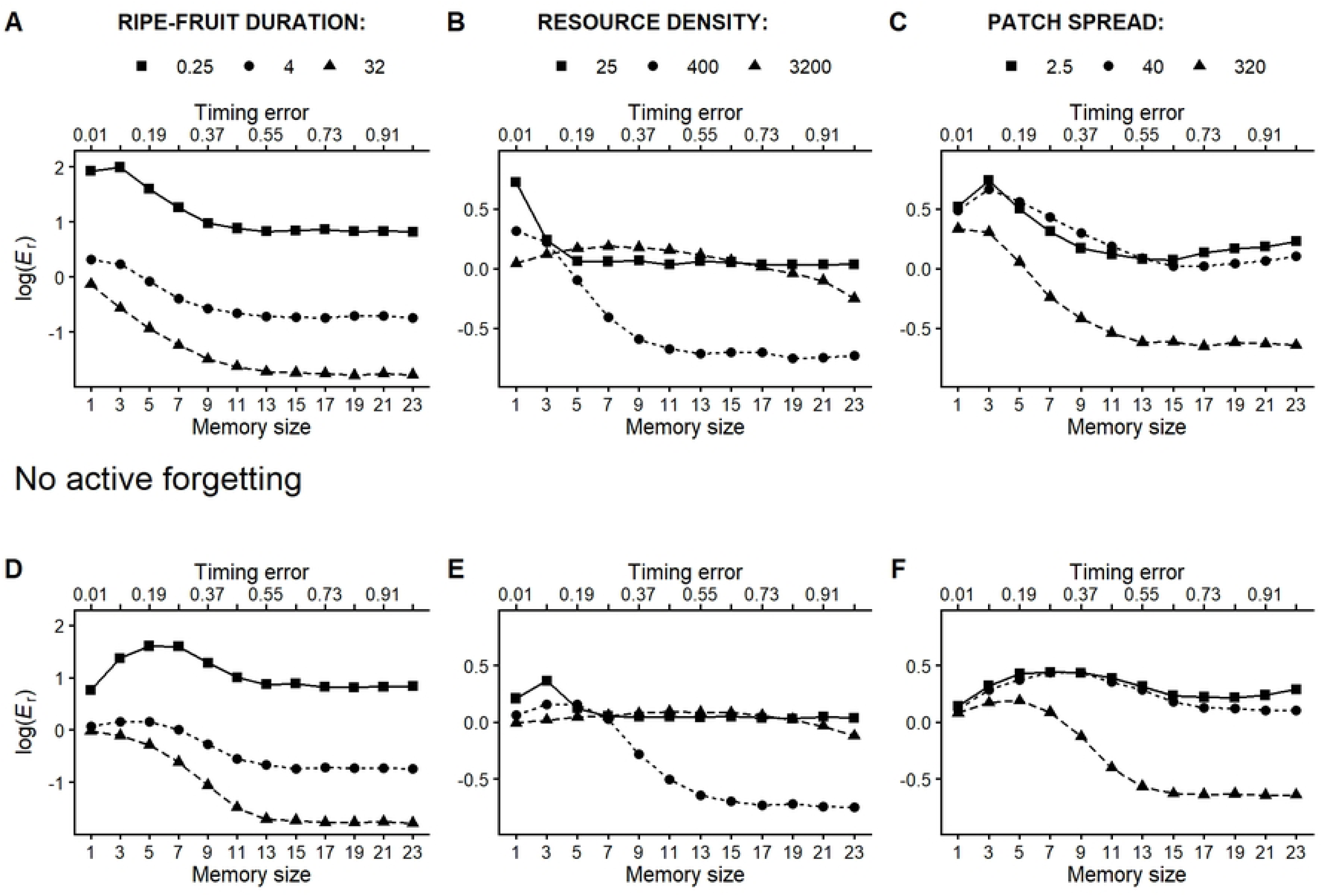
Variations in resource density, but not ripe-fruit duration and patch spread, influence the optimal quantity-quality trade-off in memory-guided foragers with (A-C) and without (D-F) the capacity to actively forget unrewarding trees. Relative foraging efficiency (log-transformed) for memory-guided agents (with active and passive forgetting abilities) with a fixed linear trade-off between the quality and quantity of memory across environments that varied in the ripe-fruit duration (A, D), productivity (B, E) and patch spread (C, F).

**Fig 6.**
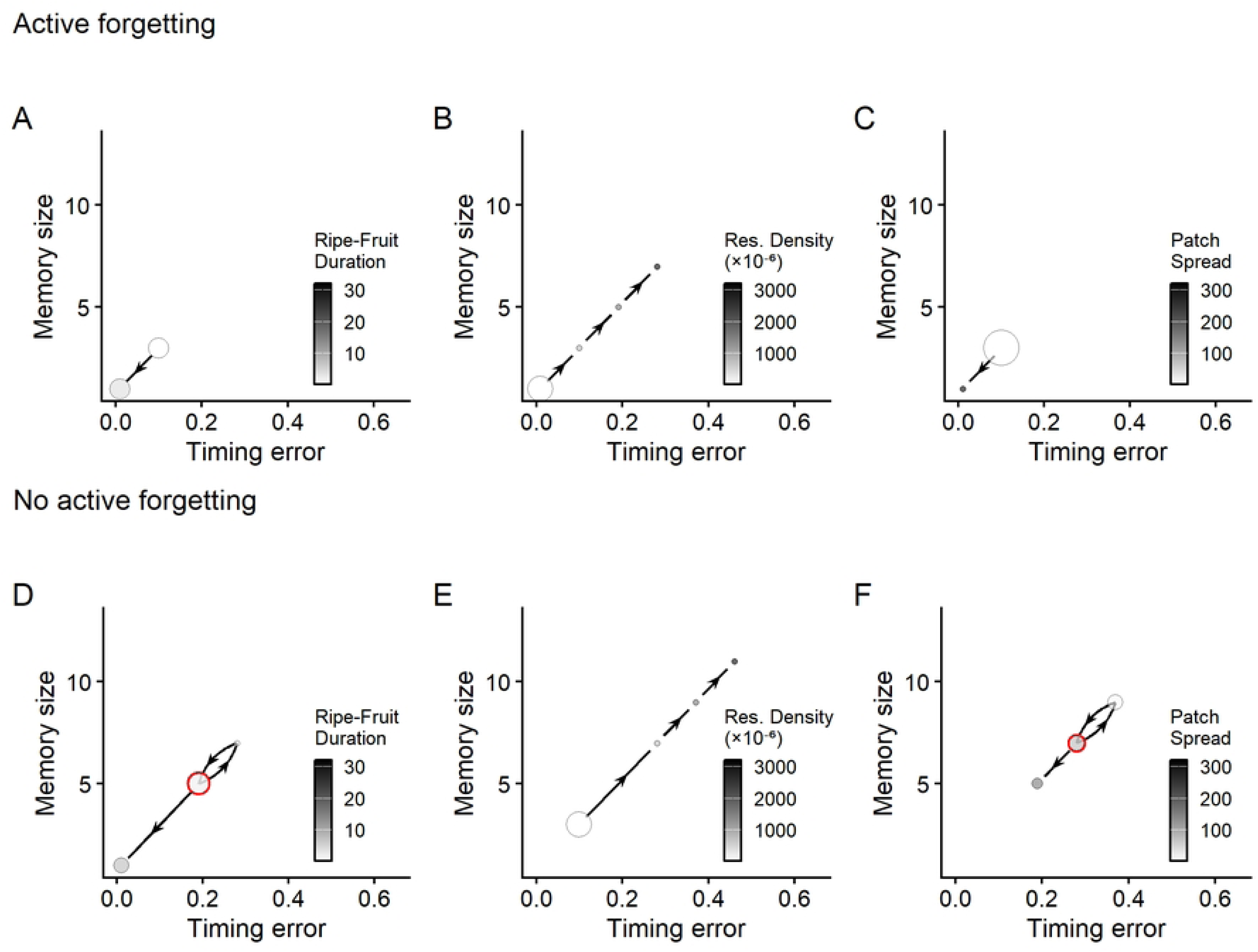
Longer ripe-fruit duration and greater patch spread shift the trade-off towards lower timing error, while increased resource density shifts it towards a larger optimal memory size. The best-performing memory parameters for memory-guided agents (with active and passive forgetting abilities) with a fixed linear trade-off between the quality and quantity of memory across environments that varied in the ripe-fruit duration (A, D), productivity (B, E) and patch spread (C, F). Each circle represents the combination of memory size and inaccuracy (i.e., quantity and quality) that yielded the highest relative efficiency in each environment. The size of the circles represent the number of environmental parameters that have that set of memory parameters as the best-performing. Circle colour corresponds to the lowest value of the environmental parameter at which that memory-parameter combination is best. The connecting lines with arrows represent the direction of change in the environmental parameter, with the arrow indicating increasing environmental value. In subplots D and F, a loop is drawn for clarity to indicate that the sequence goes back and forth once. The sequence starts at the circle with the red outline, which marks the lowest value of the environmental parameter in these subplots (ripe-fruit duration: 2.5 tu in D; patch spread: 2.5 alu in F).

With a dynamic linear and exponential trade-off (where the cost is adjusted based on the number of currently memorized trees) a similar relationship emerged as with the fixed trade-off across the different environments (Fig 5; S5 Fig).

### 3.3 Scenario 3: Active forgetting provides foraging advantages

In previous scenarios, the forager was forgetting memories both passively (i.e. fading of old memories) and actively by forgetting trees which are found empty and not containing unripe or ripe fruit. To investigate how the forgetting mechanism affects foraging efficiency under different environments and how it affects the quality-quantity trade-off, we thus repeated the simulations without such an active process where the forager only forgot old and not updated memories. In this case, a similar relationship of the quality-quantity trade-off emerged across the environmental parameters, as was found for the foragers with active forgetting abilities (Fig 5D-F). Yet, the lack of active forgetting was associated with two major absolute changes: First, a moderately larger memory size was more beneficial. This was reflected in 1) the best performing memory parameters being those with a larger memory size (S4 Fig) and 2) the optimal memory size shifting towards larger values in the quality-quantity trade-off (124% increase in optimal memory size when compared to a forager with active forgetting abilities; Fig 5; Fig 6). Second, active forgetting was associated with a higher foraging efficiency especially in ephemeral (Fig 3A; Fig A-C in S6 Fig), resource-poor environments (Fig 3B; Fig D-F in S6 Fig) and heterogeneous environments (Fig 3C; Fig G-I in S6 Fig). This foraging advantage of active forgetting was particularly exacerbated when memory size was limited (Fig C, F, I in S6 Fig).

## 4. DISCUSSION

With this study, we aimed to simulate the movement of memory-guided foragers in environments differing in the spatiotemporal distribution of resources. This enabled us to investigate how memory size, timing accuracy, and forgetting mechanisms affect foraging efficiency, and how foragers should trade off the quantity and quality of memorised information in various environments. We demonstrated that, although memorising information about food sources was generally beneficial, the extent of this benefit depended on the spatiotemporal distribution of resources. Memory conferred the greatest advantage when resources were rare, ephemeral and spatially heterogeneous. Contrary to intuition, maximising the amount and precision of information was not necessarily the most optimal strategy. Finally, despite the absence of an energetic cost of memory in our simulations, the ability to actively forget was sometimes advantageous, particularly in ephemeral, resource-poor and heterogenous environments.

### 4.1 The advantage of spatiotemporal memory

Memory-guided foraging in our simulations provided larger foraging benefits in comparison to naive foragers and this benefit depended on the resource spatiotemporal patterns. Enhanced cognitive abilities generally were most rewarding in environments with a short temporal availability of resources, and with a resource-poor and heterogeneous spatial distribution. This suggests that the utility of memory is high in environments that are characterized by intermediate to high levels of spatiotemporal complexity (Fig 3), aligning with existing theories of cognition evolution [31, 44, 55]. In more temporally stable, spatially homogeneous, and moderately productive environments, the foraging benefits of memory decline with an increasing memory size, to a point where non-memory-guided movement even can become beneficial (Fig 4). This indicates that physiological costs of acquiring and maintaining memories [56], are not the only source of limitation of memory, as has been suggested [33], since there were no direct (physiological) costs of memory in our simulations.

We observed an expected decline in the relative foraging efficiency in environments with high levels of spatial patchiness for cognitive foragers with only passive forgetting abilities [31, 44, 57]. Surprisingly, we did not observe a similar decline with a decreasing ripe-fruit duration or resource density. This likely reflects differences in available environmental information for the forager. When resources are short-lived or scarce and only with intermediate-to-high resource clustering, individuals can still effectively use memory. In such circumstances, we see that the foragers regularly encounter trees within their sensory range and collect information enabling them to anticipate ripe fruit availability. However, in highly patchy environments large areas contain few or no trees, so it becomes more difficult to build up a memory based on the scarce informative encounters with potential food sources, especially because sensory range in our model depended on tree density. As a result, the benefit of memory decreased under extreme spatial clustering when the cognitive agent lacked active forgetting abilities. In contrast, providing the simulated forager with an ability to actively forget countered this effect and at least ensured that the memories retained were more likely to be up to date. Our results thus support that the advantage of memory - in this case temporal memory specifically - is highly dependent on the distance over which foragers can detect trees that may contain fruit in the near future and the frequency with which they encounter such trees within that range [58].Together, these factors determine how much sensory information is available to guide movement. Yet, beyond an environmental effect, foraging efficiency also varied as a consequence of *what* and *how much* the forager knew.

### 4.2 The shape of quality-quantity trade-off depends on environmental conditions

Environmental structure can alter the balance between sensory information about current and future foraging opportunities. In our model, this balance determined the optimal combination of how much (quantity) is memorized and how accurately (quality) information is processed. As a result, changes in environmental structure shifted the optimal point along this quality-quantity trade-off (i.e. which memory size-accuracy combination yielded the highest foraging efficiency) in counter-intuitive ways. For example, one might expect that foragers in environments with ephemeral resources require more accurate timing abilities [16]. However, our model suggests that foragers may instead favour increased reliance on memory-based movement via an increased memory size, even at the expense of timing accuracy (Fig 5, optimum shifting to the right when ripe-fruit duration decreases; Fig 6).

#### 4.2.1 The quality-quantity trade-off affects individuals’ space use

In absence of a quality-quantity trade-off when quantity could be high at no cost of quality, we found that the timing accuracy (quality) increased foraging efficiency as it allowed more timely revisits to remembered locations (S2 Fig). However, this gain in efficiency was surprisingly modest. In contrast, memory size (quantity) had a more pronounced effect on foraging efficiency, which appears to be driven by it affecting not only the amount of remembered information but also the extent of memory-based movement. Memory-based movements solely could lead to animals becoming trapped in overexploited areas when memory use was excessive (as in [59]; S3 Fig). In our model, when no resources were detected within the sensory range, a forager with larger memory size was more likely to retain locations in memory and switch to memory based movement instead of random exploration. How it affected foraging efficiency depended strongly on both ripe-fruit duration and resource density.

#### 4.2.2 The quantity-quality trade-off is affected by the interplay between resource ephemerality and sensory acuity

In environments with less ephemeral resources (i.e. with longer ripe-fruit duration), a smaller memory size was optimal (Fig 4C; S2 Fig; S4 Fig). Here, fruits remained available longer while ripening time was fixed, thereby increasing the proportion of ripe (i.e., edible for the simulated forager) to unripe (non-edible) fruits (Fig 1D). Detection of unripe fruit is informative when combined with memory, whereas detection of ripe fruit is sufficient on its own to locate food. Consequently, in less ephemeral environments, memory use becomes less beneficial, because sensory information about currently available resources increasingly shifts the balance toward sensory, rather than memory-based, guidance.

In addition, in our model the forager prioritized sensory information about currently available food (which is reliable) over potential food that could be found by use of memory (which is likely more uncertain). In non-ephemeral environments, this implies that increasing memory size increases absolute foraging efficiency, but with diminishing returns. This stems from many remembered locations never being used because nearer ripe fruit is detected first and because there is more sensory information on currently available resources (S7 Fig). This produces negative returns in relative efficiency, because the absolute foraging efficiency of the naive forager increases more steeply under these conditions (Fig 4A–C; S7 Fig).

Conversely, when resources are more ephemeral, immediate sensory information is rarer and shorter-lived, so anticipating future opportunities becomes more important. In such environments, sampling from a larger pool of remembered locations (i.e. a larger memory size) becomes more advantageous (S4 Fig). Memory use in these ephemeral environments is beneficial even when a forager’s timing abilities are highly inaccurate (Fig 4A). This suggests that the ratio of sensory information about currently available resources versus sensory information about future opportunities is a key determinant of the value of memory. This contrasts the expectation that in ephemeral environments memory helps only when it can predict arrival time at food sources accurately [60].

#### 4.2.3 The quantity-quality trade-off is affected by resource density

In environments with fewer resources, memory became less beneficial. Yet, as opposed to the effect of ephemerality, this effect is a direct byproduct of the model construction. As resource density decreases, the detection radius also increases, as it was set to the expected nearest-neighbour distance between resources. Since there are fewer trees, it means that when moving randomly, the forager is likely to encounter a larger share of the resources in the environment (and their current status) (see Appendix D in S1 File for details). Again, this implies that based on detection only, the forager is naturally “aware” of most trees’ phenological state, making large memory sizes less beneficial.

When a trade-off between quality and quantity was introduced, the optimal combination of these two parameters was shaped by the same relationships between the memory and environmental parameters described above. Resource density, heterogeneity and the duration of ripe-fruit duration had a pronounced effect on the outcome of the quality-quantity trade-off. When resources were scarce, long-lasting or homogeneously distributed, minimal memory size (and maximal timing accuracy) was most profitable, and in resource-dense, heterogeneous and ephemeral environments this optimum shifted toward larger memory sizes (Fig 5; Fig 6). These conclusions were unchanged whether we considered the quality-quantity trade-off fixed (based on maximal memory size) or dynamic (based on realised memory size) (S5 Fig). Hereby, our model provides insight into the role of the environment on the benefits of the quantity and quality of memory, and offers a first theoretical framework for how it may shape the quality-quantity trade-off.

### 4.3 The advantage of forgetting

Memory is not only about what you know, but also what you do not know and what you forget. In particular, an active forgetting mechanism, which enabled the forager to discard unfruitful memories from its limited cognitive store, improved foraging efficiency under conditions of limited memory. Notably, active forgetting improved performance for foragers with a small memory size in environments where increasing storage capacity had no additional benefit. This suggests that the process of active forgetting does more than merely free up “space” and prevent a cognitive overload. It may enable foragers to update information [61].

Information updating may be particularly important for individuals that show site fidelity by enabling the forager to erase outdated information. Such site fidelity can be both a cause and a consequence of memory [49]. When animals restrict their space-use to the same and thus limited area, routines may emerge [48, 49]. As a result, animals can become trapped in overexploited but known areas, leading to a decrease in foraging returns, hence to an “individual-level” cost of memory. (At a population level, the cost of this restricted movement could be counterbalanced by an increase in the environmental carrying capacity for memory-guided foragers [62]).

Refreshing memories, or completely replacing them with new ones, can avoid such a memory cost. In our model, the foragers with a smaller memory size indeed were less restricted in space-use (S3 Fig). In addition, active forgetting of outdated resource locations increased foraging efficiency, especially in ephemeral, resource-poor environments, as expected from previous theoretical work [31, 33, 44]. By selectively erasing outdated information, active forgetting both promoted exploration and ensured that a limited memory capacity was mostly filled with up-to-date information about resource availability. As a consequence, the optimal memory capacity in our model was lower when active forgetting was present than when it was absent. In other words, active forgetting shortened the temporal window over which memories were retained, thereby reducing the risk of remaining in overexploited areas.

This converged with mathematical models considering a sliding-window memory in which memorized information is updated by incorporating new information that replaces the oldest memories [63]. These models likewise show that moderate memory capacities (on the order of 5–20 remembered previously consumed items) can already have a near-optimal performance [63]. By contrast, when foragers did not actively forget in our models, i.e. with only passive, time-dependent forgetting, the temporal window over which memories were retained is only defined by the fading of oldest memories. For these foragers, the optimal memory size increases, partly compensating for the slower turnover of information.

Evidently, other strategies to maintain a sufficient turnover of information exist. Foragers might also adjust the balance between exploration (in other words, information gathering) and memory-based exploitation in order to gather new and up-to-date information. For example, foragers might increase exploration in response to perceived poor local resource conditions [64] or to personal information uncertainty [65]. Such behavioural adjustment (not modelled here) could buffer the shift towards smaller optimal memory size which occurred as a response to active forgetting.

### 4.4 Concluding remarks

The benefit of cognitive capacities (memory size and accuracy) is thought to be shaped by the spatiotemporal dynamics of resources, with intermediate levels of resource predictability favouring memory the most [31, 44, 57]. Using our expertise in the natural challenges of primate foraging we here simulated a simplified, memory-guided forager to examine the effects of various distributions of resources in space, time and quantity on the foraging benefit. Our findings indicate that memory generally provided the highest benefits in environments where resources were rare and ephemeral, and moderately to highly patchy, whereas gains from memory were much smaller in temporally stable, spatially homogeneous and resource-rich environments. Our results provide only limited support for the hypothesis that memory is most beneficial in environments of intermediate spatio-temporal complexity [31]. A higher timing accuracy generally improved foraging efficiency, but increasing memory size was not always advantageous. Instead, foraging efficiency typically peaked at low to intermediate levels of memory size, with further increases leading to negative returns (and even to negative relative foraging efficiencies). For foragers with a large memory size, these negative foraging efficiencies were especially pronounced when resources were moderately abundant, non-ephemeral and homogeneously distributed. This was not due to physiological memory costs, because memory had no cost in our models. Instead, it was likely due to behavioural effects such as restricted movement and local overexploitation, which highlights a *hidden* cost of memory *per se*. These findings suggest that an effective cost of memory can emerge from how it shapes movement patterns and foraging decisions. However, a large memory size, i.e., larger brains, and improved cognitive abilities, come with increased metabolic costs [66]. While these costs were not integrated into our models, including such metabolic costs in future models could improve our understanding under which conditions enhanced cognitive abilities may be truly advantageous.

Another way in which future models could be extended concerns our conceptualisation of time tracking and errors in time estimation. In our model, this led to less accurate estimates for older memories and a slight backward telescoping effect (in which the estimated elapsed time exceeds the actual elapsed time), both of which have been observed in humans [18, 27]. An alternative extension of our model would be to systematically vary this telescoping bias [18, 19] to examine how it affects foraging efficiency and may be adaptive, thereby providing insight into the evolutionary origin of such a bias.

More generally, in our study we simulated foragers that used memory to localize food, but crucially this memory was constrained (in quality and quantity) and these constraints traded off against each other. Such constraints and trade-offs determine in what kind of environment memory and its various components are beneficial [31]. Our study takes a first step toward future models that incorporate such memory trade-offs to examine how they shape, and are shaped, by evolutionary pressures.

## Supporting Information

### S1 File. Supplementary Appendices

Appendix A. UD50. Appendix B. Dynamic trade-off. Appendix C. Relative foraging efficiency *E_f_* Appendix D. Expected fraction of encountered trees

**S1 Fig. Estimated time versus actual passed time for foragers given three different values of timing error.**

(A) Estimated time versus actual time for three individual foragers with a timing error of 0.09, 0.54 and 0.99 (black lines). The red line represents perfect time estimation, where the estimated time would be equal to the actual passed time. In this example, the actual time increment per step is set to a value of 0.2 tu (matching the time passed during random movement). (B) Mean estimated time (black line) and standard deviation (shaded area) based on 1000 simulations. The red dashed line indicates perfect time estimation.

**S2 Fig. Relative foraging efficiency (non log-transformed) of the memory-guided foragers that differed in their cognitive abilities (timing error and memory size) for various values of ripe-fruit duration (A-C), resource density (D-F) and patch spread (G-I).**

The plots in the second row (B, E, H) and third row (C, F, I) show cross-sections of the contour plots (A, D, G) for different values of memory size and timing error, respectively. The red lines indicate the threshold where there is no relative foraging benefit compared to a naive forager.

**S3 Fig. The area (in alu^2^) of the utilization distribution under the 50% isopleth (UD50) for memory-guided foragers across environments that varied in ripe-fruit duration (A), resource density (B) and patch spread (C).**

Memory size corresponds to the number of memorised trees. Points correspond to the average UD50 across 300 simulations.

**S4 Fig. The best-performing memory parameters of the two types of foragers across environments which varied in ripe-fruit duration (A), resource density (B) and patch spread (C).**

Each circle represents the combination of memory size and inaccuracy (i.e., quantity and quality) that yielded the highest relative efficiency in each environment. The size of the circles represent the number of environmental parameters that have that set of memory parameters as the best-performing. The circle with the red outline indicates the environmental parameter with the lowest environmental value (i.e., ripe-fruit duration: 2.5 tu [A], resource density: 25 x 10⁻⁶ trees/alu² [B] and patch spread: 2.5 alu [C]). The connecting lines with arrows represent the direction of the change in the environmental parameter, with the arrow indicating increasing environmental value.

**S5 Fig. Relative foraging efficiency (log-transformed) for memory-guided foragers (with active forgetting abilities) with a dynamic linear trade-off between the quality and quantity of memory.**

The dynamic trade-off was modelled as a linear (A-C) and an exponential relationship (D-F) and was simulated across environments that varied in the duration of ripe-fruit duration (A, D), resource density (B, E) and patch spread (C, F).

**S6 Fig. Foraging efficiency of memory-guided foragers with active forgetting abilities relative to the efficiency of memory-guided foragers that did not possess the ability for active forgetting in different environments and with different combinations of memory parameters (timing error and memory size).**

Memory size corresponds to the number of memorised trees. Environments differed in ripe-fruit duration (A-C), resource density (D-F) and patch spread (G-I). The plots in the second row (B, E, H) and third row (C, F, I) show cross-sections of the contour plots (A, D, G) for different values of memory size and timing error, respectively. The red lines indicate the threshold where there is no relative foraging benefit compared to a forager with only a passive forgetting mechanism. See Appendix C in S1 File for the calculation of Relative foraging efficiency *E_f_*.

**S7 Fig. Absolute efficiency of the best-performing memory-guided foragers and naive foragers across environments that varied in ripe-fruit duration (A), resource density (B) and patch spread (C).**

Points correspond to the average absolute foraging efficiency across 300 simulations.

**S1 Table. Ranges of values for key parameters that were used in the simulations.**

Values in bold were the default values.

